# Coat color mismatch improves survival of a keystone boreal herbivore: energetic advantages exceed lost camouflage

**DOI:** 10.1101/2021.09.24.461654

**Authors:** Joanie L. Kennah, Michael J. L. Peers, Eric Vander Wal, Yasmine N. Majchrzak, Allyson K. Menzies, Emily K. Studd, Rudy Boonstra, Murray M. Humphries, Thomas S. Jung, Alice J. Kenney, Charles J. Krebs, Stan Boutin

## Abstract

Climate warming is causing asynchronies between animal phenology and environments. Mismatched traits, like coat color change mismatched with snow, can decrease survival. However, coat change does not serve a singular adaptive benefit of camouflage, and alternate coat change functions may confer advantages that supersede mismatch costs. We found that mismatch reduced rather than increased, autumn mortality risk of snowshoe hares in Yukon by 86.5 %. We suggest that the increased coat insulation and lower metabolic rates of winter acclimatized hares confer energetic advantages to white mismatched hares that reduce their mortality risk. We found that white mismatched hares forage 17-77 minutes less per day than matched brown hares between 0 and -10 °C, thus lowering their predation risk and increasing survival. We found no effect of mismatch on spring mortality risk, where mismatch occurred at warmer temperatures, suggesting a potential temperature limit where the costs of conspicuousness outweigh energetic benefits.

## Introduction

Phenological mismatch is one of the most documented pathways by which climate change negatively impacts species (Radchuk et al. 2019, Visser and Gienapp 2019). Earlier onset of spring and delayed onset of winter have the potential to cause incongruous timing of seasonal phenotypes (Møller et al. 2008, Lehikoinen 2011, Kudo and Ida 2013). Mismatch occurs in the timing of numerous seasonal traits such as calving date with plant growth onset, and laying date with peak of key food sources, and has resulted in reduced reproductive success and recruitment (Post and Forchhammer 2008, Reed et al. 2013). However, the costs associated with phenological mismatches vary within species across populations (Heard et al. 2012, Doi et al. 2017). Species are often adapted to broad ranges of ecological conditions, particularly those with large geographic distributions (Valladares et al. 2014). Local adaptations and variable selection pressures across environmental gradients alter the magnitude of phenological mismatch across populations (Phillimore et al. 2010, Gordo and Doi 2012, Porkert et al. 2014). Such spatial variability in phenology across ecological conditions may also involve differences in the mechanistic pathways governing the demographic costs and benefits associated with phenological mismatch across species ranges.

An example of phenological mismatch that occurs in species across multiple taxa is coat and plumage color change mismatched with snow onset and melt (Zimova et al. 2016, Pedersen et al. 2017, Atmeh et al. 2018, Melin et al. 2020). At least 21 bird and mammal species in the Northern Hemisphere change color biannually and improved camouflage is considered the primary function of this change (Mills et al. 2018, Zimova et al. 2018) As snow cover duration is forecasted to decrease across the Northern Hemisphere (Danco et al. 2016), coat and plumage color mismatch is likely to increase. Mismatch may reduce survival due to decreased camouflage (Atmeh et al., 2018; Zimova et al., 2016; Melin et al., 2020). However, aside from color change, high-latitude species benefit from other winter acclimatization strategies meant to increase cold tolerance and endure periods of food shortage, including increasing insulation, decreasing lower critical temperature, altering activity patterns, and, ultimately reducing daily energy requirements (Humphries et al. 2005, Fuglesteg et al. 2006, Sheriff et al. 2009b). Accordingly, coat color transitions coincide with multi-trait change that differentiates long photoperiod, i.e., summer, from short photoperiod, i.e., winter, phenotypes (Lovegrove 2005, Boratyński et al. 2016). As such, the thermal and energetic benefits provided by a more insulative, white coat and associated metabolic and thermoregulatory adaptations may outweigh the negative costs of color mismatch in colder conditions.

The snowshoe hare (*Lepus americanus*) is a keystone species distributed across the boreal forests of North America (Krebs et al. 1995) that undergoes seasonal coat color change to match the presence of snow (Ferreira et al. 2017). The initiation of coat color change in snowshoe hares is likely affected by photoperiod (Nagorsen 1983) and in the absence of evolutionary change, is predicted to become increasingly mismatched with anticipated reductions in snow cover duration (Brown and Mote 2009, Mills et al. 2013). Coat color mismatch may impact snowshoe hare demography, as recent studies have reported high mortality rates in mismatched snowshoe hares at multiple locations in the southern extent of their range, presumably due to increased conspicuousness to predators (Zimova et al., 2014; Wilson et al., 2018). However, the thermal benefits of winter acclimatization in hares, including reduced metabolic rate (Sheriff et al. 2009a), may also affect susceptibility to predation and ultimately survival.

White winter-acclimatized snowshoe hares benefit from lower energetic demands compared to brown summer-acclimatized hares. Indeed, while temperatures below 0 °C increase energetic requirements for summer hares, white winter hares remain in their thermoneutral zone until temperatures below -10 °C (Sheriff et al. 2009a). As such, lower energetic demands reduce foraging requirements for winter-acclimatized hares (Balluffi-Fry *et al*., In Review). Balancing the trade-off between obtaining sufficient food to meet energetic requirements and avoiding predators is a central assumption of prey behavior theory (McNamara and Houston 1987, Lima and Dill 1990). Therefore, white mismatched hares may benefit from lower energetic requirements, reduced foraging time, and thus reduced predator exposure. These benefits could compensate for the adverse effects of conspicuousness, particularly when seasonal temperatures remain low and the energetic demands for brown summer acclimatized hares are elevated (Balluffi-Fry *et al*., In Review). Geographic variation in winter adaptations and acclimatization exists across the broad geographic range of the snowshoe hare (Sheriff et al. 2009b, Gigliotti et al. 2017). As such, the effects of coat color mismatch may vary across populations according to the relative importance of the reduced camouflage cost relative to energy conservation benefits in different ecological contexts.

Here, we test the hypothesis that reduced foraging requirements with winter acclimatization reduces the costs of coat color mismatch in snowshoe hares. To examine this, we monitored the survival, coat color, and foraging time of individuals over the autumn and spring in southwest Yukon, Canada. First, we predict that mismatched white hares will spend less time foraging than matched brown individuals, particularly below the thermoneutral zone of summer brown hares (i.e. 0 °C; Sheriff *et al*. 2009a). If this foraging difference and thus reduced time spent vulnerable to predation outweighs the costs of conspicuousness, we further predict no difference in survival between matched and mismatched individuals. However, if camouflage loss is the primary driver of predation risk during coat color change, regardless of foraging differences, we expect that mismatched hares are more likely to be predated than camouflaged individuals, echoing results from previous studies in the southern extent of their range (Zimova et al. 2016, Wilson et al. 2018). We found that white mismatched snowshoe hares experiencing cold temperatures in snowless environments benefitted from reduced foraging time and thus increased survival relative to brown matched hares.

## Methods

### Study area

We studied snowshoe hares for three autumns (September 1^st^ to December 1^st^ of 2015, 2016, and 2017) and four springs (March 1^st^ to May 31^st^ of 2015, 2016, 2017, and 2018) in southwestern Yukon, Canada (Lat: 60.9 N, Long: -138.0 W). Snowshoe hares have been monitored for over 40 years in this region (Krebs et al., 2018). Our study area consists predominantly of white spruce (*Picea glauca*), trembling aspen (*Populus tremuloides*) and balsam poplar (*Populus balsamifera*). Gray willow (*Salix glauca*) and dwarf birch (*Betula glandulosa*) dominate the understory. The main predators of snowshoe hares in this region include Canada lynx (*Lynx canadensis*), coyotes (*Canis latrans*), goshawks (*Accipiter gentilis*), and great horned owls (*Bubo virginianus*) (Peers et al. 2020). Snowshoe hares went through the increase, peak, and early decline phase of their population cycle during our study period (Krebs et al. 2018).

### Field methods

The study area was divided into three 35-ha snowshoe hare trapping areas, located within ∼ 8 km of each other (Peers et al. 2020). We captured snowshoe hares using Tomahawk live-traps (Tomahawk Live Trap Co. Tomahawk, WI, USA) baited with alfalfa and rabbit chow. Traps were set 30 minutes before sunset and checked either three hours after sunset or at sunrise. We attached a numbered ear tag to each hare to identify individuals on subsequent recaptures, and we assessed coat-color during each capture. To evaluate coat color, we examined hares from the front and sides and visually estimated their percentage white coat to the nearest 5%. We later binned coat color in 10% white categories for analyses to account for inter- and intra- observer ranking variability. We consider 10% bins as reasonably precise given that intra- and inter- observer intraclass correlation coefficients (ICC) for coat color assessment were high (ICC>0.9 in all cases, See Appendix S1: Table S1). To monitor survival, we fit hares weighing > 1100g (n=347) with very high frequency (VHF) collars that were each equipped with a mortality sensor (Model SOM2380, Wildlife Materials Inc., USA, or Model MI-2M, Holohil, Canada, both < 27 ± 1 g). We performed mortality checks of VHF collared hares almost daily, i.e., 96.3% of checks occurred within 1 to 3 days. To monitor behavior, we also fit a subset (*n* =102) of VHF collared hares with an accelerometer (model Axy3, 4 g, Technosmart, Rome, Italy). Accelerometers measure force variation on three different axes and are increasingly being used to infer behavior in free-ranging animals (Mikkelsen et al. 2019, Studd et al. 2019). Fully equipped collars with both VHF and accelerometers had a total weight below 2.5% of each individual’s body mass. Handling and collaring procedures were approved by the University of Alberta Animal Care and Use Committee (Protocol: AUP00001973).

We measured snow depth, snow cover, and temperature throughout our study period. We measured snow depth on >60% of days at three locations per trapping area, in relatively open forest, to the nearest 0.5 cm. Days with missing snow depth records were linearly interpolated using the “zoo” function in the zoo package in R (Zeileis et al. 2021). We measured snow cover by visually assessing daily landscape photographs from three camera traps installed on each trapping area. We calculated a combined average daily snow cover value to the nearest 10% in our study region. We converted % snow cover to a binary type variable above or below 60% snow cover (presence/absence) for the autumn seasons, as there were very few instances when snow cover estimates were between 0% and 100%. We measured temperature at least six times a day on each trapping area using a minimum of 2 temperature loggers (ibutton, DS1922L, Maxim Integrated, Whitewater, USA) to obtain a single average daily temperature value for each trapping area.

### Measuring coat color mismatch

Coat color mismatch was defined as the difference between hare percent white (10% bins) and the daily percent snow cover (10% bins for both autumn and spring). For all analyses, we treated mismatch as a binary variable, defining mismatch as greater than 50% difference between hare % white and snow cover (%). As such, mismatched hares were white (> 50 % white) individuals in a snowless (< 50% snow cover) environment. Considering that brown mismatched hares in a snowy environment were rare (1% of trapping records), we did not consider this type of mismatch in analyses. Although the threshold for mismatch used in some previous studies is 60% contrast (Mills et al. 2013, Wilson et al. 2018), mismatch at this contrast threshold was rare in our study region, i.e., in 11% of trapping records, so we used 50% as our mismatch threshold to increase our sample size. That being said, analyses using 40% or 60% thresholds for mismatch revealed similar results (Appendix S1: Tables S5, S6, S9, S10).

### Effect of coat color mismatch on survival

To evaluate the effect of coat color mismatch on snowshoe hare survival, we generated Cox’s proportional hazards (CPH) models (Cox and Oakes 1984) with the “coxph” function in the survival package in R (Therneau et al. 2021). The CPH model is a semi-parametric approach used to analyze binary response data, in our case: alive or dead (Sievert and Keith 1985). We monitored 347 hares and recorded 41 deaths over four springs and 34 deaths over three autumns. We excluded mortality checks that exceeded seven days to limit the uncertainty in the timing of death events (Murray and Bastille-Rousseau 2020). We censored 15 individuals whose collars were removed before the end of the study period and six individuals with permanently missing VHF signals. We pooled data from different years, trapping areas, and sex, as exploratory analysis indicated that none of those variables had a significant effect on autumn or spring mortality risk (Appendix S1: Table S2). Considering that coat color was assessed only during capture opportunities (on average every 13.1 ± SD: 10.8 days per individual), we assigned coat color for each record in our survival analysis as the nearest coat color assessment completed in the field (average difference of 4.95 ± SD: 3.70 days between telemetry check and coat-color assessment). We removed telemetry records where a coat color assessment within 14 days did not exist to ensure that coat color and derived mismatch values were an accurate representation of each individual at the time of the telemetry check. Results from models using survival records within 8 days of a coat-color assessment were qualitatively similar to those we obtained with our chosen 14-day threshold (Appendix S1: Table S3).

We generated three competing CPH models for both autumn and spring. The first model included snow cover and snow depth, based on prior evidence of snow effects on hare survival (Meslow and Keith 1971, Peers et al. 2020). Our second model included those same snow variables in addition to coat color mismatch, our variable of interest. The third model was the null (intercept-only) model. We used Akaike Information Criterion for our model selection (Akaike 1974) and identified our top model based on AIC_c_ (Burnham and Anderson 2002) with the package AICcmodavg (Mazerolle 2019). We assessed multicollinearity in our top model using the variance inflation factor (VIF) and ensured no variables had VIF’s greater than 2. The proportionality assumption of CPH models, which implies that the hazard ratio (HR; i.e., risk of death) is assumed to be constant over time (Joshua Chen and Liu 2006), was met for our top spring and autumn CPH model. Our results were not affected by informative censoring, as we found qualitatively similar results for both spring and autumn model coefficients when we treated censored individuals as deaths (Murray and Bastille-Rousseau 2020) (Appendix S1: Table S4).

### Effect of coat color mismatch on time spent foraging

To test our proposed mechanistic pathway, whereby white mismatched hares experience reduced energetic requirements leading to reduced foraging time (Balluffi-Fry *et al*. In Review; Sheriff *et al*. 2009a), we used linear mixed-effects models using the “lmer” function in the package lme4 (Bates et al. 2015). Daily time spent foraging (minutes) was derived from tri-axial accelerometer data using behavioral classifications previously developed in this hare population (see Studd et al., 2019 for more information on classification methods). Daily time spent foraging was classified over 4 second intervals at a 96% accuracy (Studd et al. 2019). We recorded 1505 daily foraging records from 66 hares over the three autumns and 838 daily foraging records from 44 hares over the four springs. Similar to our survival analysis, we only kept foraging records that were within 14 days of a coat-color assessment (average difference of 4.48 ± 3.51 (SD) days). We reran our top foraging time models with data restricted to daily foraging records that were within 8 days of a coat-color assessments instead to ensure that our results were not affected by this 14-day threshold, and obtained qualitatively similar results (Appendix S1: Table S8). To eliminate the potential of seasonal changes in foraging impacting our results (Griffin et al. 2005), we restricted our data to only the autumn and spring periods when snow cover was ≤ 50%, i.e., mismatch was possible given our chosen threshold and therefore both matched and mismatched individuals occurred simultaneously.

We generated four linear mixed-effects models per season to test for differences in daily minutes spent foraging (our response for all models) between matched brown hares and mismatched white hares and their responses to changes in temperature. We included a random effect for individual ID in all models to control for non-independence of data. We included sex as a fixed factor in all spring models only, as exploratory data analysis indicated that sex had a significant effect on time spent foraging for spring but not autumn (Appendix S1: Table S7). Furthermore, we included year as a fixed effect in each model to account for potential effects of yearly changes in predation risk on hare foraging behavior (Shiratsuru et al. 2021). Our first model included two fixed effects, temperature and year. Our second model included temperature, year, and coat color mismatch, and our third model included the same variables as the second in addition to an interaction between mismatch and temperature. Our fourth model was a null intercept-only model. We checked model fit using marginal and conditional R-squared calculated using the “r.squaredGLMM” function in the package MuMIn (Barton 2020), according to Nakagawa *et al*. 2017. We used Akaike Information Criterion (Akaike 1974) to rank our four competing models and identified our top model in each season based on AIC_c_ (Burnham and Anderson 2002).We completed all statistical analyses in R version 3.6.2 (2019) (R Core Team, 2019). We considered results where P ≤0.05 as significant and reported all means with ± 1 standard error.

## Results

Permanent snow cover date, i.e., 100% snow cover without melting until the spring, was variable across our autumn seasons, occurring almost 3 weeks later in 2015 (November 3^rd^) than in 2016 (October 16^th^) and 2017 (October 17^th^). Completion of snowmelt date, i.e., no more snow on ground, was similar across study years (May 6^th^, 2015, May 1^st^ 2016, May 2^nd^ 2017 and May 1^st^ 2018). When considering both seasons and all years together, the prevalence of coat color mismatched hares that contrasted with their snowless environment was low (14% of trapping records) in our population. Mismatch occurred more frequently in the autumn (19% of trapping records) than the spring (8% of trapping records). The autumn with the latest permanent snow cover arrival date, i.e., 2015, had the highest prevalence of mismatch (33% of records). Prevalence of mismatch in the autumns of 2016 and 2017 were 10% and 13% of trapping records, respectively. Spring mismatch was consistent across years around 10% (2015-9% of trapping records, 2016-10%, 2018-12%), with the exception of 2017 when only 1% of trapped hares were mismatched.

### Effect of coat color mismatch on mortality

The CPH model with the strongest support in both seasons included snow depth, snow cover and mismatch (Appendix S1: Table S11, S12 & S13). However, the second highest ranking CPH model for spring, i.e., the model including only snow variables, was within 2 ΔAICc (AICc = 0.09) from our top spring CPH model (Appendix S1: Table S11). Mortality risk for mismatched hares in autumn was significantly reduced (*z*= -2.43; P=0.02) relative to matched hares (Hazard Ratio (HR)= 0.135; 95% Confidence Intervals (CI): 0.027, 0.679; Fig. 1a). In contrast, coat color mismatch was positively correlated with mortality risk for hares in the spring (Fig. 1b), but this effect was non-significant (*z*= 1.60; P= 0.11). Models were qualitatively similar regardless of our classification of mismatch, except when considering mismatch as a minimum 40% contrast between coat color and snow cover; in this case mismatch significantly increased mortality risk in the spring (HR= 6.780; 95% CI: 2.390, 19.240; *z*= 3.60; P<0.001). Snow depth (*z*= -2.29; P= 0.02) and snow cover (*z*= 2.98; P=0.003) significantly affected mortality risk in the top spring model, but not in the top autumn model. In spring, the risk of dying decreased as snow depth increased (HR=0.95; 95% CI: 0.92, 0.993; Appendix S1: Fig S1a) and mortality risk increased as snow cover increased (HR=1.046; 95% CI: 1.01, 1.08; Appendix S1: Fig S1b).

**Fig 1.**
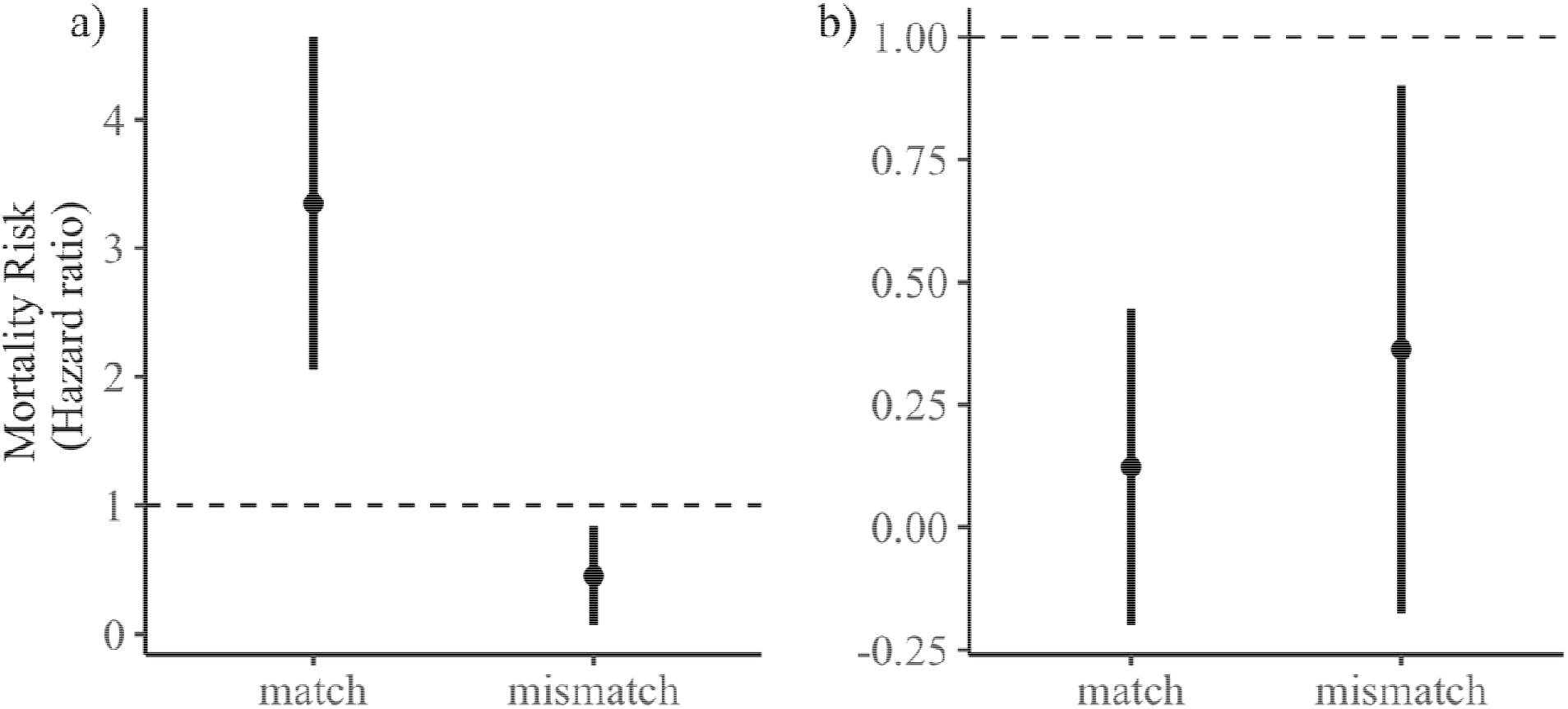
The modelled effect of coat color mismatch on snowshoe hare mortality risk, generated from our top supported CPH model for a) autumn and b) spring. Points represent predicted hazard ratios (HR) for matched and mismatched hares when snow depth and snow cover are held at zero. Error bars represent predicted standard errors, and the dashed line represents baseline mortality risk (i.e., HR=1).

### Effect of coat color mismatch on foraging time

Across our study years, hares foraged on average 706 ± 2.29 minutes per day in the spring and 751 ± 1.65 minutes per day in the autumn. Coat color mismatch was an important predictor of daily foraging time in the autumn, but not the spring (Appendix S1: Table S14 and S15). The top model for autumn foraging time included coat color mismatch, temperature, year, and the interaction between temperature and mismatch (Table 1). As autumn temperature decreased, mismatched hares decreased daily foraging time, whereas matched hares increased foraging time (Fig. 2a; Table 1). For instance, when the temperature was - 8 °C, brown-matched hares foraged 65 minutes more per day than white-mismatched hares (Fig. 2a). The top model for spring included temperature, year, and sex (Table 1), When coat color mismatch was included in our spring foraging models, its effect on daily foraging time was non-significant (*t* =-0.759, *P*>0.05).

**Table 1.**
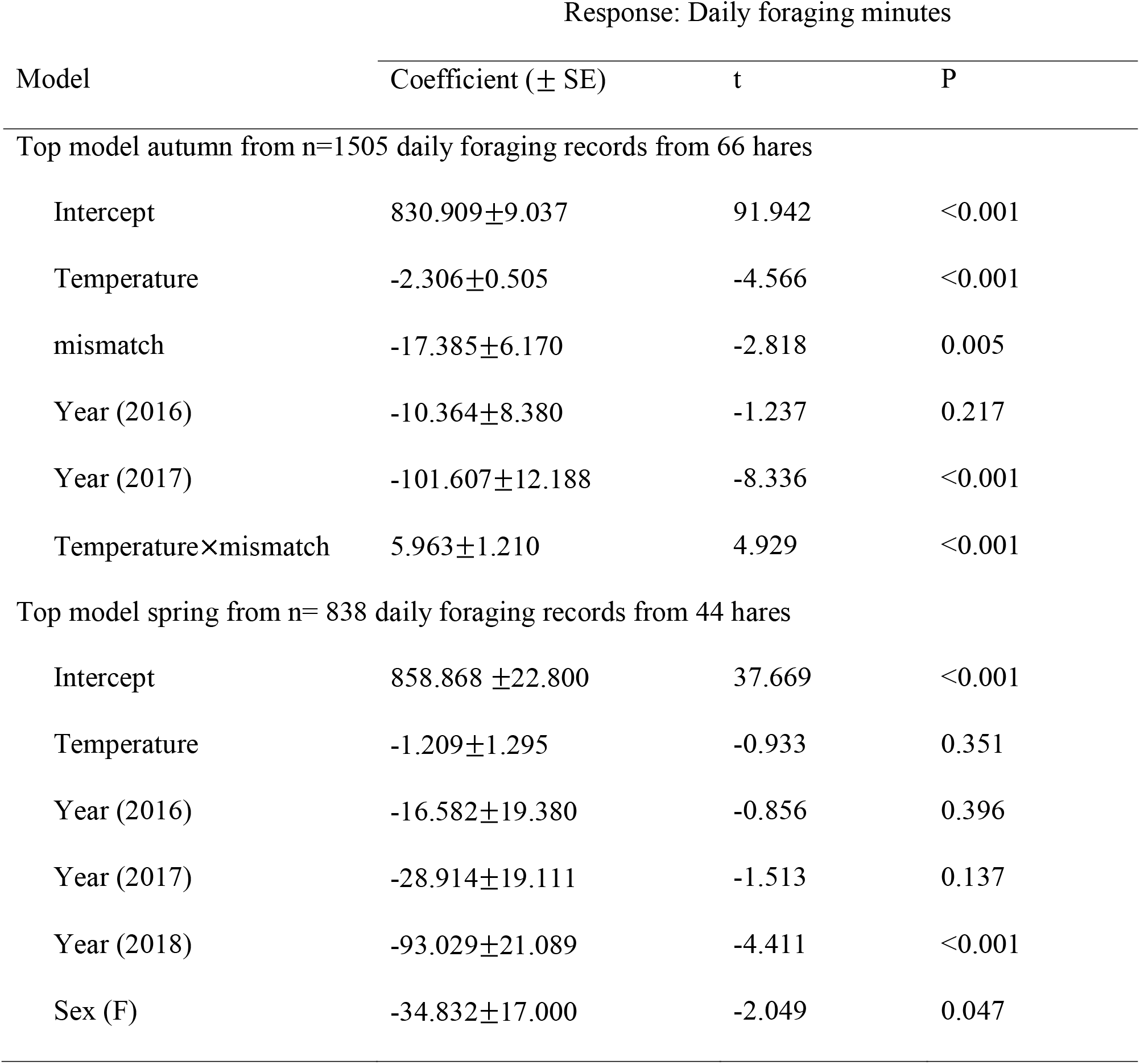
Summary of variables included in top-ranking linear mixed-effects daily foraging time models for snow-free autumn and spring periods. Daily foraging time was considered in minutes. Both autumn and spring models also include individual ID as a random effect and the spring model includes sex as a random effect.

| Model | Response: Daily foraging minutes |  |  |
| --- | --- | --- | --- |
| | Coefficient (± SE) | t | P |
| Top model autumn from n=1505 daily foraging records from 66 hares |  |  |  |
| Intercept | 830.909±9.037 | 91.942 | <0.001 |
| Temperature | -2.306±0.505 | -4.566 | <0.001 |
| mismatch | -17.385±6.170 | -2.818 | 0.005 |
| Year (2016) | -10.364±8.380 | -1.237 | 0.217 |
| Year (2017) | -101.607±12.188 | -8.336 | <0.001 |
| Temperature×mismatch | 5.963±1.210 | 4.929 | <0.001 |
| Top model spring from n= 838 daily foraging records from 44 hares |  |  |  |
| Intercept | 858.868 ±22.800 | 37.669 | <0.001 |
| Temperature | -1.209±1.295 | -0.933 | 0.351 |
| Year (2016) | -16.582±19.380 | -0.856 | 0.396 |
| Year (2017) | -28.914±19.111 | -1.513 | 0.137 |
| Year (2018) | -93.029±21.089 | -4.411 | <0.001 |
| Sex (F) | -34.832±17.000 | -2.049 | 0.047 |

**Fig 2.**
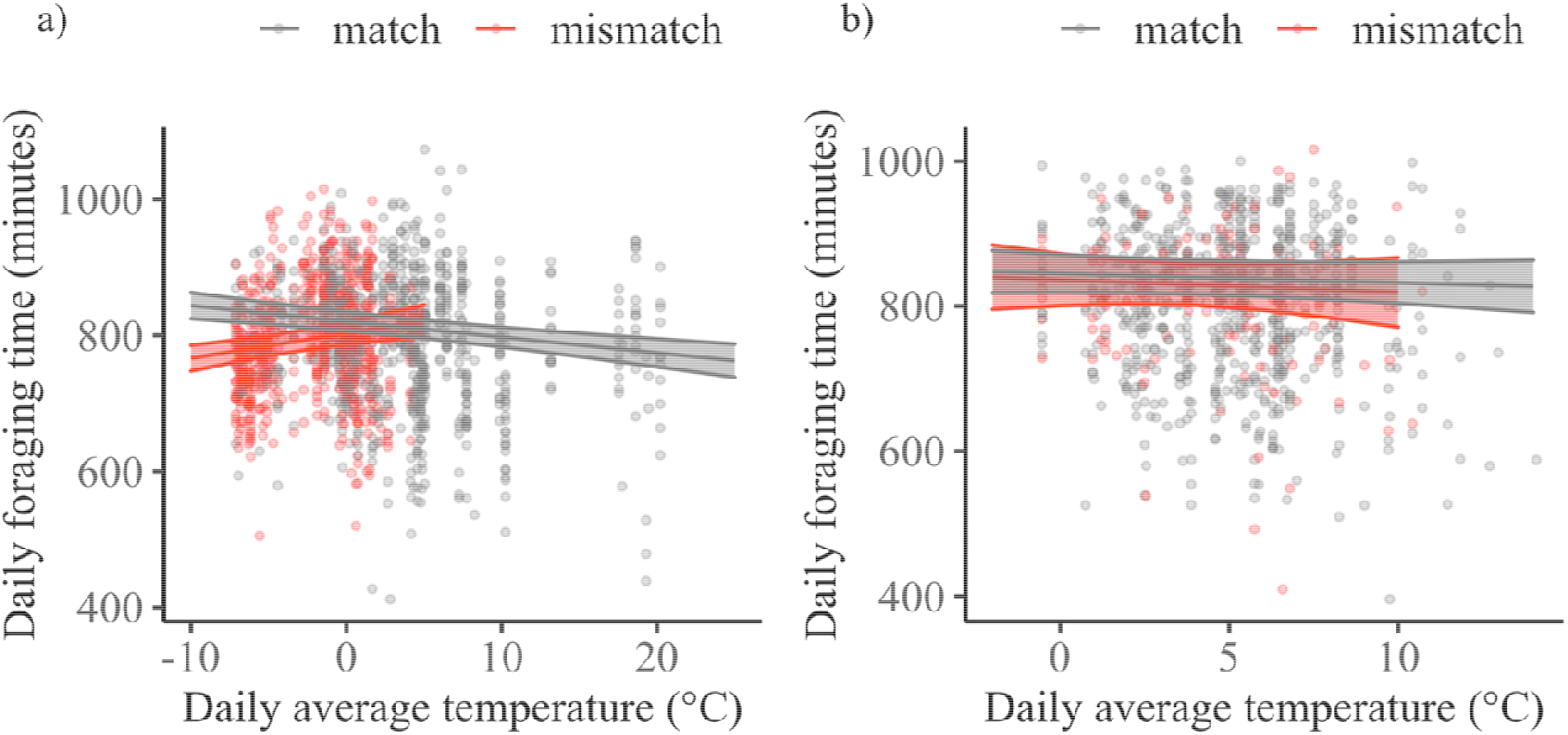
Modelled effect of temperature on daily foraging time (minutes) for matched and mismatched snowshoe hares in the snow-free period of a) autumn (marginal R^2^= 0.12, conditional R^2^=0.32) and b) spring (marginal R^2^=0.13, conditional R^2^=0.28) of 2016 (the year with the most data). Data points show daily foraging records for individuals across all study years and predicted foraging time of mismatched hares is restricted to temperatures where mismatched hares occurred in our study. Predicted values for daily spring foraging time are for males.

## Discussion

Phenotypes and climate change can vary widely within a species’ distribution, as can phenological mismatch and its consequences on survival. Elucidating potential unifying mechanisms is crucial to reconcile varied responses to phenological mismatch. We evaluated the effect of coat color mismatch on snowshoe hare survival in a northern population and further tested a potential mechanism that may influence this effect. We hypothesized that the thermal and energetic benefits of winter acclimatization in white hares, i.e., increased coat insulation and reduced metabolic rate (Sheriff et al. 2009a, Gigliotti et al. 2017), ultimately reduce their foraging requirements (Balluffi-Fry *et al*. In Review) and thus predation risk, which may influence the costs of coat color mismatch. Surprisingly, we found that mismatched hares had a higher survival than matched hares in the autumn (Fig. 1a) but that survival did not differ between matched and mismatched hares in the spring (Fig. 1b). Although this result contradicts previous studies that link coat color mismatch in snowshoe hares to reduced survival (Zimova et al. 2016, Wilson et al. 2018), our proposed mechanism for why this might be the case is supported. Mismatched white hares spent significantly less time foraging than matched individuals in the autumn (Fig. 2a), presumably due to the thermal and energetic benefits of winter acclimatization. Indeed, reduced foraging time likely decreases exposure to predators and subsequently improves survival (Fig 1a). We reconcile our findings with those of previous studies with a unifying factor: temperature.

Matched hares foraged longer than mismatched white individuals in the autumn, and this difference was pronounced at lower (< –3°C) temperatures (Fig. 1a). Given the wide range of ecological contexts, selection pressures, and local adaptations that exist across the distribution of snowshoe hares (Gigliotti et al. 2017), the cost-benefit ratio of lost camouflage versus energy conservation may vary across populations experiencing different temperatures. For example, northern populations experiencing cold temperatures benefit from the energetic advantages of winter coats despite mismatch during snow-free periods, whereas southerly populations experiencing warmer temperatures may not. Indeed, adverse survival effects associated with mismatch in southern snowshoe hare populations in Montana (Zimova et al. 2016) and Wisconsin (Wilson et al. 2018) occur in regions that experience warmer temperatures than those in southwestern Yukon (Fig. 2). During the period when mismatch is possible in Montana, autumn temperatures can range from ∼ 3°C to 17 °C and spring temperatures can range from ∼ 4°C to 20 °C.

The seasonal differences in mismatch effects on survival and foraging time that we found within our study population highlight temperature as a unifying factor affecting the survival costs of coat color mismatch. In spring, mismatch did not influence mortality risk (Fig. 1b) and matched and mismatched hares spent similar amounts of time foraging (Fig. 2b). Mismatched hares in the spring occurred at temperatures (−0.5 °C to 11°C, Fig. 2b) that were approximately within the thermoneutral zone of both summer and winter-acclimatized hares (Sheriff et al. 2009a). In contrast, mismatched hares in the autumn experienced temperatures between -7°C and 4°C (Fig. 2a) which fall below the lower critical temperature for summer-acclimatized brown hares, but not winter-acclimatized white hares (Sheriff et al. 2009a). Animals must increase their energetic expenditure when they are exposed to temperatures outside of their thermoneutral zone (Kingma et al. 2012), which may represent a likely mechanism explaining the longer foraging time in matched brown hares in the autumn relative to mismatched white hares (Fig. 2a). These results further support that the thermal and energetic benefits of winter acclimatization may outweigh the costs of coat color mismatch at cold temperatures.

Although camouflage is thought to be the primary adaptive benefit of coat color polymorphism, like most traits, alternate benefits, e.g., thermal and physiological, exist (Caro 2005, Duarte et al. 2017, Zimova et al. 2018). We found that these alternate benefits offset the costs of camouflage loss at cold temperatures. Our proposed hypothesis, whereby the thermal and energetic benefits of winter acclimatization may influence coat color mismatch effects through reduced time spent foraging, has the potential to reconcile intraspecific variation among other snowshoe hare populations and merits testing in other color changing species, i.e. arctic hares (*Lepus arcticus*), mountain hares (*Lepus timidus*). Climate change-induced variation in temperature and precipitation regimes are likely to vary across species ranges (Loarie et al. 2009). Such variation in climate change effects will be particularly large for species with broad distributions, i.e., circumboreal color-changing species. Ultimately, as temperatures in the Northern Hemisphere are projected to warm (Danco et al. 2016), northern snowshoe hare populations are likely to reach the threshold (> –3°C) at which the energetic benefits of white coats are lost, and survival costs driven by coat color mismatch could occur (Zimova et al. 2016, Wilson et al. 2018). However, elucidating the mechanisms through which phenological mismatches may be operating is essential to enable predictions on broad-scale changes in species distributions.

## Acknowledgements

We thank the numerous field technicians who worked on this project, as well as members of the Wildlife Evolutionary Ecology lab at Memorial University of Newfoundland for comments on earlier versions of this manuscript. We thank Sean Konkolics and Alec Robitaille for assistance and help with statistical analyses and R code. We also thank A. MacDonald and her family for long-term access to her trapline. We thank the Champagne and Aishihik First Nations, and Kluane First Nation, for allowing this work within their traditional territory. This work was supported by the Natural Sciences and Engineering Research Council of Canada, Northern Studies Training Program, the University of Alberta Northern Research Award programme, the Association of Canadian Universities for Northern Studies, the Wildlife Conservation Society Canada, the W. Garfield Weston Foundation, Government of Yukon, and Earth Rangers.

