## Supplementary material for "Coat color mismatch improves survival of a keystone boreal herbivore: energetic advantages exceed lost camouflage": Supplemementary Information

Journal: Ecology

Supplementary Information

Appendix S1 – Additional Results………………………………………………………..pg. 2-17

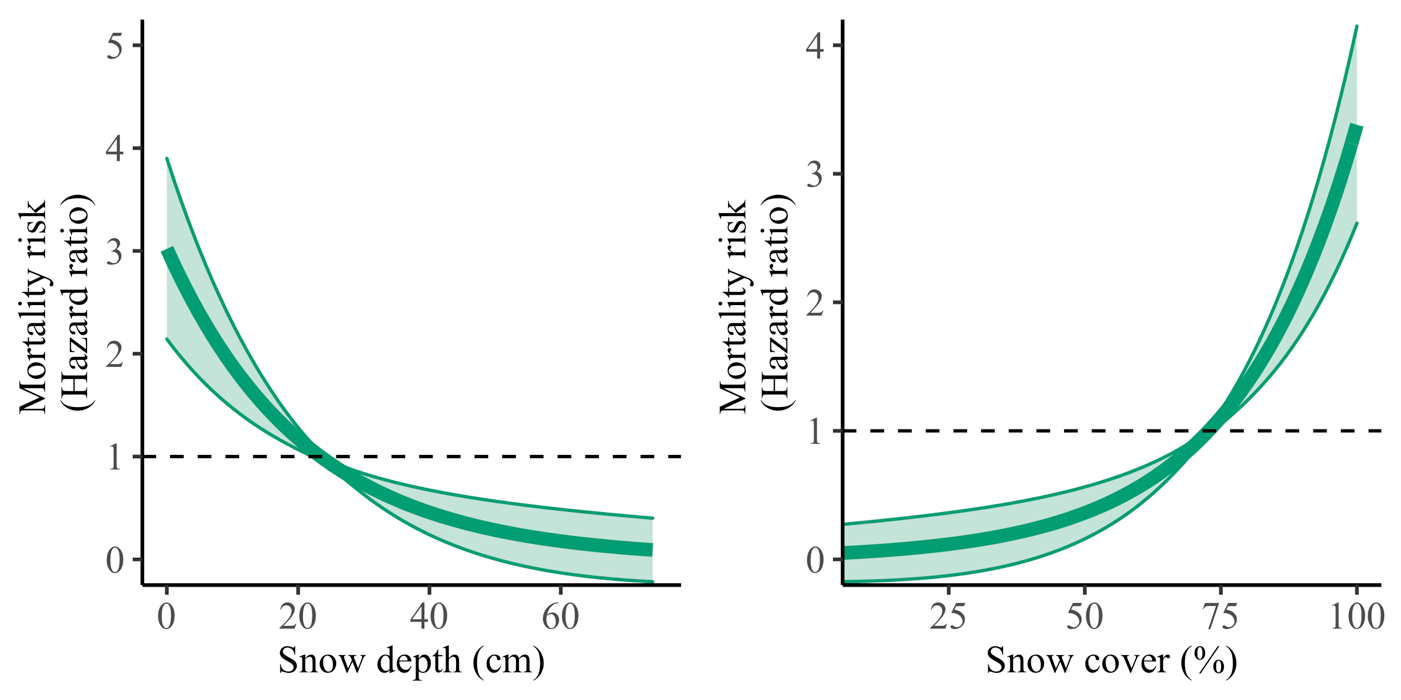

a)	b)

Fig S1. Effect of a) snow depth (snow cover held constant at its mean) and b) snow cover (snow depth held constant at its mean) on matched snowshoe hare spring mortality risk. Generated from n=41 deaths for 229 hares recorded over 4 spring seasons (2015-2018). The shaded area represents standard error. The dashed lines represent baseline hazard ratio (HR) (1), HR>1 indicates increased risk of dying and HR<1 indicates reduced risk of dying.

Table S1. Intraclass correlation coefficients (ICC) describing intra-observer repeatability and inter-observer repeatability of measurements taken within and between three observers. Each observer ranked coat colour three times to the nearest 5% for 50 hares (total of n=150 rankings for each observer). Inter-rater repeatability was estimated using the average of the three measurements for each observer.

|  | ICC | 95% confidence intervals | F (df1, df2) | p-value |
| --- | --- | --- | --- | --- |
| Observer 1 | 0.978 | 0.966, 0.987 | 137 (49,100) | < 0.001 |
| Observer 2 | 0.957 | 0.933, 0.974 | 67.7 (49,100) | < 0.001 |
| Observer 3 | 0.949 | 0.92, 0.969 | 56.9 (49,100) | < 0.001 |
| Between observers | 0.913 | 0.855, 0.947 | 32.6 (49,100) | < 0.001 |

Table S2. Effect of various predictors that may affect spring and autumn hazard ratio (HR). Each variable was tested in a univariate model during exploratory data analysis to evaluate its effect on HR.

| Variable | Levels | Season | | | | HR | | | | z | | | | P |
| --- | --- | --- | --- | --- | --- | --- | --- | --- | --- | --- | --- | --- | --- | --- |
| Sex | Reference=Male | | | |  | | | |  | | | |  | |
|  | Female | Autumn | | | | 1.465 | | | | 1.04 | | | | 0.298 |
|  |  | Spring | | | | 0.929 | | | | -0.23 | | | | 0.818 |
| Trapping area | Reference=TA-1 | | |  | | | |  | | | |  | | |
|  | TA-2 | Autumn | | | | 0.702 | | | | -0.914 | | | | 0.361 |
|  |  | Spring | | | | 0.556 | | | | -1.632 | | | | 0.103 |
|  | TA-3 | Autumn | | | | 0.415 | | | | -1.867 | | | | 0.062 |
|  |  | Spring | | | | 0.675 | | | | -0.930 | | | | 0.352 |
| Year | Reference= 2015 | |  | | | |  | | | |  | | | |
|  | 2016 | Autumn | | | | 0.519 | | | | -1.670 | | | | 0.095 |
|  |  | Spring | | | | 0.532 | | | | -1.381 | | | | 0.167 |
|  | 2017 | Autumn | | | | 1.091 | | | | 0.179 | | | | 0.858 |
|  |  | Spring | | | | 1.247 | | | | 0.539 | | | | 0.590 |
|  | 2018 | Spring | | | | 0.830 | | | | -0.366 | | | | 0.714 |

Table S3. Hazard ratios (HR) and 95% confidence intervals generated for each variable of top CPH models when only survival records occurring within 8 days of known coat colour assessment are considered. Values that are bolded represent significant effects and italicized values represent P values < 0.1. HR>1 indicates increased risk of dying and HR<1 indicates reduced risk of dying.

| Variable | Spring top model:  HR~SD+SC+mm | Autumn top model:  HR~SD+SC+mm |
| --- | --- | --- |
| Snow depth (SD) | 0.960 (0.914, 1.010) | *0.831* (0.685, 1.009) |
| Mismatch (mm;factor) | 3.004 (0.335, 26.962) | **0.128** (0.023, 0.707) |
| Snow cover (SC) | **1.064** (1.016, 1.114) | 1.113(0.073, 16.966) |

Table S4. Hazard ratios (HR) and 95% confidence intervals generated for each variable of top CPH models when missing individuals are treated as deaths. Values that are bolded represent significant effects. HR>1 indicates increased risk of dying and HR<1 indicates reduced risk of dying.

| Variable | Spring top model:  HR~SD+SC+mm | Autumn top model:  HR~SD+SC+mm |
| --- | --- | --- |
| Snow depth (SD) | **0.960** (0.926, 0.996) | **0.890** (0.798, 0.992) |
| Mismatch (mm;factor) | 2.284 (0.608, 8.580) | **0.134** (0.027, 0.675) |
| Snow cover (SC) | **1.035** (1.009, 1.062) | 0.810 (0.074, 8.893) |

Table S5. Hazard ratios (HR) and 95% confidence intervals generated for each variable of top autumn CPH model when mismatch is defined as hares at least 40% whiter than their environment (environmental whiteness measured by snow cover). Values that are bolded represent significant effects and italicized values represent P values < 0.1. HR>1 indicates increased risk of dying and HR<1 indicates reduced risk of dying.

| Variable | Spring top model:  HR~SD+SC+mm | Autumn top model:  HR~SD+SC+mm |
| --- | --- | --- |
| Snow depth (SD) | *0.953* (0.915, 0.993) | *0.904* (0.809, 1.010) |
| Mismatch (mm:factor) | *6.780* (2.390, 19.240) | *0.285* (0.073, 1.107) |
| Snow cover (SC) | *1.056* (1.025, 1.087) | 1.376 (0.138, 13.734) |

Table S6. Hazard ratios (HR) and 95% confidence intervals generated for each variable of top autumn CPH model when mismatch is defined as hares at least 60% whiter than their environment (environmental whiteness measured by snow cover). Values that are bolded represent significant effects and italicized values represent P values < 0.1. HR>1 indicates increased risk of dying and HR<1 indicates reduced risk of dying.

| Variable | Spring top model:  HR~SD+SC+mm | Autumn top model:  HR~SD+SC+mm |
| --- | --- | --- |
| Snow depth (SD) | **0.953** (0.915, 0.993) | *0.903* (0.809, 1.009) |
| Mismatch (mm:factor) | 3.640 (0.770, 17.207) | **0.111** (0.019, 0.657) |
| Snow cover (SC) | **1.044** (1.014, 1.075) | 0.776 (0.077, 7.814) |

Table S7. Effect of sex on spring and autumn daily foraging minutes when tested in a linear mixed-effects model including sex (fixed), and individual ID as a random factor.

| Variable | Levels | Season | | Coefficient ($\pm$ SE) | | t | | P |
| --- | --- | --- | --- | --- | --- | --- | --- | --- |
| Sex | Reference=Male | |  | |  | |  | |
|  | Female | Autumn | | -22.09 | | -1.56 | | 0.124 |
|  |  | Spring | | -45.25 | | -2.632 | | <0.001 |

Table S8. Summary of top-ranking autumn and spring linear mixed-effects daily foraging time models during snow-free seasons when the maximum number of days elapsed between coat colour assessment and corresponding foraging records is eight days.

|  | Response: Daily foraging minutes | | |
| --- | --- | --- | --- |
| Model | Coefficient ($\pm$ SE) | t | P |
| Top autumn model from n=1382 daily foraging records from 65 hares | | | |
| Intercept | 834.122$\pm$9.176 | 90.905 | <0.001 |
| Temperature | -2.285$\pm$0.516 | -4.428 | <0.001 |
| mismatch | -17.899$\pm$6.183 | -2.895 | 0.004 |
| Year (2016) | -10.799$\pm$8.495 | -1.271 | 0.204 |
| Year (2017) | -110.748$\pm$13.181 | -8.402 | <0.001 |
| Temperature$\times$mismatch | 6.257$\pm$1.221 | 5.125 | <0.001 |
| Top spring model from n= 791 daily foraging records from 43 hares | | | |
| Intercept | 857.642 $\pm$22.946 | 37.377 | <0.001 |
| Temperature | -1.542$\pm$1.297 | -1.189 | 0.235 |
| Year (2016) | -13.325$\pm$19.571 | -0.681 | 0.499 |
| Year (2017) | -26.519$\pm$19.286 | -1.375 | 0.175 |
| Year (2018) | -89.545$\pm$21.118 | -4.240 | <0.001 |
| Sex (F) | -30.925$\pm$17.247 | -1.793 | 0.080 |

Table S9. Summary of top-ranking autumn and spring linear mixed-effects daily foraging time models during snow-free seasons when mismatch is defined as hares at least 40% whiter than their environment (environmental whiteness measured by snow cover.

|  | Response: Daily foraging minutes | | |
| --- | --- | --- | --- |
| Model | Coefficient ($\pm$ SE) | t | P |
| Top autumn model from n=1505 daily foraging records from 66 hares | | | |
| Intercept | 831.650$\pm$8.981 | 92.603 | <0.001 |
| Temperature | -2.405$\pm$0.533 | -4.511 | <0.001 |
| mismatch | -17.968$\pm$5.812 | -3.091 | 0.002 |
| Year (2016) | -9.604$\pm$8.046 | -1.194 | 0.233 |
| Year (2017) | -103.287$\pm$12.173 | -8.485 | <0.001 |
| Temperature$\times$mismatch | 5.549$\pm$1.071 | 5.181 | <0.001 |
| Top spring model from n=838 daily foraging records from 44 hares | | | |
| Intercept | 858.868$\pm$22.800 | 37.669 | <0.001 |
| Temperature | -1.209$\pm$1.295 | -0.933 | 0.351 |
| Year (2016) | -16.582$\pm$19.380 | -0.856 | 0.396 |
| Year (2017) | -28.914$\pm$19.111 | -1.513 | 0.137 |
| Year (2018) | -93.029$\pm$21.089 | -4.411 | <0.001 |
| Sex(F) | -34.832$\pm$17.000 | -2.049 | 0.047 |

Table S10. Summary of top-ranking autumn and spring linear mixed-effects daily foraging time models during snow-free seasons when mismatch is defined as hares at least 60% whiter than their environment (environmental whiteness measured by snow cover.

|  | Response: Daily foraging minutes | | |
| --- | --- | --- | --- |
| Model | Coefficient ($\pm$ SE) | t | P |
| Top autumn model from n=1505 daily foraging records from 66 hares | | | |
| Intercept | 828.614$\pm$8.917 | 92.926 | <0.001 |
| Temperature | -2.183$\pm$0.494 | -4.420 | <0.001 |
| mismatch | -15.691$\pm$6.419 | -2.445 | 0.015 |
| Year (2016) | -9.339$\pm$8.408 | -1.111 | 0.267 |
| Year (2017) | -100.086$\pm$12.148 | -8.239 | <0.001 |
| Temperature$\times$mismatch | 6.065$\pm$1.260 | 4.813 | <0.001 |
| Top spring model from n= 838 daily foraging records from 44 hares | | | |
| Intercept | 858.868 $\pm$22.800 | 37.669 | <0.001 |
| Temperature | -1.209$\pm$1.295 | -0.933 | 0.351 |
| Year (2016) | -16.582$\pm$19.380 | -0.856 | 0.396 |
| Year (2017) | -28.914$\pm$19.111 | -1.513 | 0.137 |
| Year (2018) | -93.029$\pm$21.089 | -4.411 | <0.001 |
| Sex (F) | -34.832$\pm$17.000 | -2.049 | 0.047 |

Table S11. Selection of CPH models predicting spring snowshoe hare hazard ratio (HR) (risk of dying) generated from n= 41 deaths from a total of 229 hares recorded over 4 spring seasons (2015-2018), considering number of parameters (k) and ranked from most to least support by ∆AIC, Akaike model weights (w) and Log Likelihood (LL).

| Model | k | ∆AIC_c_ | w | LL |
| --- | --- | --- | --- | --- |
| HR~Snow depth+Snow cover +mismatch | 3 | 0.00 | 0.50 | -191.12 |
| HR~Snow depth+Snow cover | 2 | 0.09 | 0.48 | -192.17 |
| HR~1 (null) | 0 | 5.72 | 0.03 | -196.98 |

Table S12. Selection of CPH models predicting autumn snowshoe hare hazard ratio (HR) (risk of dying) generated from n= 34 deaths from a total of 218 hares recorded over 3 autumn seasons (2015-2018), considering number of parameters (k) and ranked from most to least support by ∆AIC, Akaike model weights (w) and Log Likelihood (LL).

| Model | k | ∆AIC_c_ | w | LL |
| --- | --- | --- | --- | --- |
| HR~Snow depth+Snow cover+mismatch | 3 | 0.00 | 0.89 | -157.78 |
| HR~1 (null) | 0 | 5.43 | 0.06 | -161.50 |
| HR~Snow depth+Snow cover | 2 | 5.59 | 0.05 | -163.58 |

Table S13. Hazard ratios (HR) and 95% confidence intervals generated for each variable of top CPH models. Values that are bolded represent significant effects and italicized values represent P values < 0.1. HR>1 indicates increased risk of dying and HR<1 indicates reduced risk of dying.

| Variable | Spring top model:  HR~SD+SC+mm | Autumn top model:  HR~SD+SC+mm |
| --- | --- | --- |
| Snow depth (SD) | **0.953** (0.916, 0.993) | *0.903* (0.808, 1.009) |
| Mismatch (mm;factor) | 2.940 (0.781, 11.062) | **0.135** (0.027, 0.679) |
| Snow cover (SC) | **1.046** (1.015, 1.077) | 0.726 (0.065, 8.084) |

Table S14. Selection of mixed linear models predicting daily foraging minutes for snowshoe hares in snow-free autumn seasons generated from N=1505 daily foraging records from 66 hares over 3 years (2015-2017). Considering number of parameters (k) and ranked from most to least support by ∆AIC, Akaike model weights (w) and Log likelihood (LL). . Marginal R^2^ (Mar R^2^) and conditional R^2^ (Cond R^2^) are included to show each model’s fit. Response variable for all models is daily foraging time in minutes (Forage_T). Fixed effect predictors include: temperature (temp), mismatch (mm) and year (yr), and individual ID (ID) is included as a random factor .

| Model | k | ∆AIC_c_ | w | LL | Mar. R^2^ | Con. R^2^ |
| --- | --- | --- | --- | --- | --- | --- |
| Forage_T~temp*mm+yr+ID | 8 | 0.00 | 1 | -8659.89 | 0.12 | 0.32 |
| Forage_T~temp+mm+yr+ID | 7 | 22.08 | 0 | -8671.95 | 0.10 | 0.31 |
| Forage_T~temp+yr+ID | 6 | 47.99 | 0 | -8685.91 | 0.09 | 0.29 |
| Forage_T~1+ID | 3 | 103.65 | 0 | -8716.76 | 0 | 0.32 |

Table S15. Selection of mixed linear models predicting daily foraging minutes for snowshoe hares in snow-free spring seasons generated from N= 838 daily foraging records from 44 hares over 4 years (2015-2018). Considering number of parameters (k) and ranked from most to least support by ∆AIC, Akaike model weights (w) and Log likelihood (LL). Marginal R^2^ (Mar R^2^) and conditional R^2^ (Cond R^2^) are included to show each model’s fit. Response variable for all models is daily foraging time in minutes (Forage_T). Fixed effect predictors include: temperature (temp), mismatch (mm) and year(yr), and individual ID (ID) is included as a random factor .

| Model | k | ∆AIC_c_ | w | LL | Mar. R^2^ | Con. R^2^ |
| --- | --- | --- | --- | --- | --- | --- |
| Forage_T~temp+yr+sex+ (1\|ID) | 8 | 0.00 | 0.57 | -4947.70 | 0.12 | 0.28 |
| Forage_T~temp+mm+yr+sex+(1\|ID) | 9 | 1.16 | 0.32 | -4947.26 | 0.13 | 0.28 |
| Forage_T~temp*mm+yr+sex+(1\|ID) | 10 | 3.19 | 0.13 | -4947.25 | 0.13 | 0.28 |
| Forage_T~1+(1\|ID) | 3 | 20.26 | 0 | -4962.90 | 0 | 0.28 |
